# Partial Face Visibility and Facial Cognition: An Electroencephalography and Eye-Tracking Investigation

**DOI:** 10.1101/2023.09.07.556282

**Authors:** Ingon Chanpornpakdi, Yodchanan Wongsawat, Toshihisa Tanaka

## Abstract

Face masks became a part of everyday life in the SARS-CoV-2 pandemic. Previous studies showed that the face cognition mechanism involves holistic face processing, and the absence of face features could lower cognition ability. This is opposed to the experience during the pandemic, when people were able to correctly recognize faces, although the mask covered a part of the face. This paper shows a strong correlation in face cognition based on the EEG and eye-tracking data between the full and partial faces. We observed two event-related potentials, P3a in the frontal lobe and P3b in the parietal lobe, as subcomponents of P300. Both P3a and P3b were lowered when the eyes were invisible, and P3a evoked by the nose covered was larger than the full face. The eye-tracking data showed that 16 out of 18 participants focused on the eyes associated with the EEG results. Our results demonstrate that the eyes are the most crucial feature of facial cognition. Moreover, the face with the nose covered might enhance cognition ability due to the visual working memory capacity. Our experiment shows the possibility of people recognizing faces using both holistic face processing and structural face processing. Furthermore, people can recognize the masked face as well as the full face in similar cognition patterns due to the high correlation in the cognition mechanism.

## 1 Introduction

The spread of the SARS-CoV-2 pandemic significantly affected our lifestyle, necessitating the mandatory use of masks. Before the pandemic, the face cognition mechanism of holistic face processing was the focus. The holistic processing of faces leads to an enhanced ability to identify faces and enhances visual short-term and long-term memory for face recognition purposes (Tanaka and Farah, 1993; Curby and Gauthier, 2007; Curby et al, 2009). Covering a part of the face reduces face cognition and matching performance abilities (Tanaka and Farah, 1993; Nguyen and Pezdek, 2017; Carragher and Hancock, 2020). Throughout the experience of the pandemic spread, it highlighted that a human can still perceive faces correctly even if a mask covers parts of the face. Hence, the questions of how humans recognize a partially covered face and whether holistic processing still works in this face cognition mechanism are raised.

To study face cognition mechanisms in visual perception studies, the rapid serial visual presentation (RSVP) paradigm is commonly used (Lees et al, 2018). Rapid Serial Visual Presentation (RSVP) is a type of visual stimulus presentation where the target and non-target images are presented rapidly, with the target image presented less frequently. During the stimulus presentation, brain activity can be recorded using neuroimaging methods such as non-invasive methods, electroencephalography (EEG), functional magnetic resonance imaging (fMRI), magnetoencephalography (MEG), functional near-infrared (fNIR), and invasive methods such as electrocorticography (ECoG). Our study focuses on EEG response as it directly measures brain electrical activity and provides efficient information about brain activity owing to its high temporal resolution. The recording procedure is also simple, and the machine is portable compared to other methods (Nicolas-Alonso and Gomez-Gil, 2012; Ramadan and Vasilakos, 2017).

The EEG response observed from the RSVP task is an event-related potential (ERP). This ERP is evoked due to a cognition mechanism when a participant perceives the stimulus. Therefore, ERP is a temporal waveform that occurs in a fixed time window following the onset of a stimulus (Sur and Sinha, 2009; Woodman, 2010). It contains several components in one response. Each component is named according to the polarity of the peak and the latency when the peak was induced.

In face cognition, several ERP components corresponding to face images are induced. Allison et al (1994) first found that faces evoked a negative peak at a latency of approximately 192 msec (N200). Subsequently, Bentin et al (1996) found that face images evoked a negative peak at 172 msec (N170) in the posterior temporal area absent when a non-face image was shown. Additionally, P190 has a similar property to the vertex positive potential (VPP) of latency 150–200 msec in the occipitotemporal cortex (Jeffreys and Tukmachi, 1992) and can be regenerated from a single face-responsive neuron (Bentin et al, 1996). N170 and VPP have been shown to have similar response properties and are considered face-sensitive ERP components (Joyce and Rossion, 2005; Eimer, 2011; Cai et al, 2013). However, N200 and P300 potentials are more relevant than N170 and VPP potentials for target and non-target face cognition (Eimer, 2011; Cai et al, 2013). P300 is a positive peak observed around 250–700 msec after the stimulus was perceived (Lees et al, 2018). The P300 consists of two components: a faster P3a reflecting attention focus and a slower P3b associated with memory storage (Polich and Criado, 2006; Polich, 2007). The P300 component is typically observed to have a larger amplitude around the parietal region (Farwell and Donchin, 1988; Hruby and Marsalek, 2002). It is known to be associated with target cognition and is commonly used as a biomarker in visual perception studies.

When it comes to the study of partial face, covering the eye component has also delayed the N170 latency but did not affect the N170 amplitude, which implies that the N170 component did not primarily reflect the brain response to the eye component (Bentin et al, 1996; Eimer, 1998). Similarly, Nemrodov et al (2014) performed a task on identifying upright and inverted faces and reported that N170 was delayed when the eyes were absent, whereas the amplitude was unaffected. Eimer (2000) also studied the removal of external features (i.e., the shape of the face, ears, neck, and hair) and internal features (i.e., eyes, nose, and mouth) in face cognition. N170 was reduced and delayed, meaning that N170 might be responsible for the structural encoding of the face as a whole rather than for encoding a part of the face. Żochowska et al (2022) investigated the self-face, familiar faces, and unfamiliar faces with and without the mask and found that the N170 of the faces with the surgical-like mask has longer latency than the full face. In contrast to the idea of holistic face processing in the aforementioned studies, they found that elicited P300 was stronger in the surgical-like face than the full face. In addition, Susac et al (2004) experimented on the emotion and inverted face with only sunglasses on the target face. They found that the N170 and P300 were elicited with a stronger amplitude (more negative for N170 and more positive for P3) for the target with sunglasses covering the eyes. This result could be due to either the absence of the eye component or the identification easiness of a target with glasses among full faces with neutral and smiling facial emotions. Based on the previous studies, the question of whether humans recognize partial faces as holistic face processing and the effect of the visibility of each face component and their combinations on face cognition remains unclear. Hence, our study aims to clarify these questions about the partial face.

In our previous study, we studied partial face cognition using machine learning to classify the target and non-target images based on ERP evoked (Chanpornpakdi and Tanaka, 2023b). Our results showed that the presence of the eyes affected the accuracy of the model, and we assumed that eyes are the most crucial component in face cognition. However, more investigation is needed on the characteristic ERP pattern and the eye gaze during the task to conclude our assumption. Therefore, we introduced eye-tracking in this study.

The eye movements, such as rapid saccades, prolonged fixations, and systematic scanning, can convey a variety of cognition functions (Lewis and Krupenye, 2022), also known as the eye-mind hypothesis (Just and Carpenter, 1980). Based on this hypothesis, many researchers implement the eye tracker to investigate cognitive function. Kano and Tomonaga (2009) found that humans and chimpanzees have similar behavior in eye gaze. Both humans and chimpanzees are spotted on animals or humans rather than the background and the face rather than the body in the natural behavior task. Kano and Tomonaga (2010) found that humans and chimpanzees focused on facial features by looking at the eyes first, then the mouth. Chakravarthula et al (2021) found that people focus on the eyes during the face identification task and focus on the mouth during the ethnic group categorization task. Since the pandemic, researchers also have eyes on partial face cognition, performed an experiment using an eye tracker on the masked face cognition task, and found that the mask reduced cognitive ability (Hsiao et al, 2022).

Thus, in this paper, we hypothesize that 1) the ERP evoked in full face and partial face with the eyes visible is stronger than that of the face without the eyes presented and 2) the participants focus on the eye component the most. We used EEG combined with eye-tracking face cognition by comparing the full face to the partial face cognition ability. This helps us understand how the human brain works and associates with the eye gaze. Seven conditions of the full and partial face were used as stimuli. All the conditions were full face, face with eyes covered, face with nose covered, face with mouth covered, face with eyes and nose covered, face with eyes and mouth covered, and face with nose and mouth covered.

## 2 Methods

### Participants

The experiment was performed on 31 healthy participants without neurological disorders, but only the data from 19 participants (average age of 28.21 ± 2.34, range of 23 – 33 years, nine females) were used for further analysis due to the noise contamination in EEG data and low gaze sample (lower than 85%), low accuracy, and low sensitivity in the eye-tracking data. All the participants voluntarily participated in the experiment and had normal or corrected-to-normal eyesight. This study was approved by the Mahidol University Central Institutional Review Board (COA No. MU-CIRB 2016/198.1511). All the participants provided their written informed consent before the experiment and received compensation for participating.

### Experimental Design

The experiment was designed with the same face image stimuli and similar experimental flow in the no-button press task of our previous studies (Chanpornpakdi and Tanaka, 2023a,b). The face images used in the experiment were from a public dataset of the “Ethnic Origins of Beauty (Les origines de la beauté)” project by Ivanova N. (available at lesoriginesdelabeaute.com) (Ivanova, n.d.). Images were resized to 710 × 555 pixels and cropped to a visual angle of 7.4 × 5.2 c/d, showing only the face area. The experiment consisted of the training task and the main task. Each task contains seven blocks with seven face conditions: full face, face with eyes covered, face with nose covered, face with mouth covered, face with eyes and nose covered, face with eyes and mouth covered, and face with nose and mouth covered. (see Chanpornpakdi and Tanaka (2023a,b) for more details)

### Training Task

This task allowed the participants to learn the target face, which was an unfamiliar face image to all the participants. One designated Asian target face and two non-target Asian faces were used in this experiment. The stimulus presentation was made using the Psychophysics Toolbox Version 3 (Brainard, 1997) in MATLAB 2020a, showing each face ten times/condition (30 trials in total/face condition). The image was presented until the participant pressed a button to answer whether or not the presented image was the target image. If the answer was correct, the beep sounds at 800, 1,300, and 2,000 Hz were played consecutively. If the answer was wrong, only a beep sound at 800 was played.

### Main Task

The target face was trained in the training task, and five Asian faces not used in the training task were used as stimulus images. We created the stimulus to be presented using Tobii pro lab version 1.194 (Tobii, Stockholm, Sweden). Figure 1 shows the experimental flow of the main task, containing seven blocks of seven face conditions. For each block, the face images were presented 100 times each for 200 msec with a fixation of 500 msec between the stimuli to prevent the attentional blink (Raymond et al, 1992). The participants were asked to count the number of target faces shown during each block and reported the number counted after each block. EEG, eye gaze, and the number of target faces recognized were recorded.

**Fig. 1.**
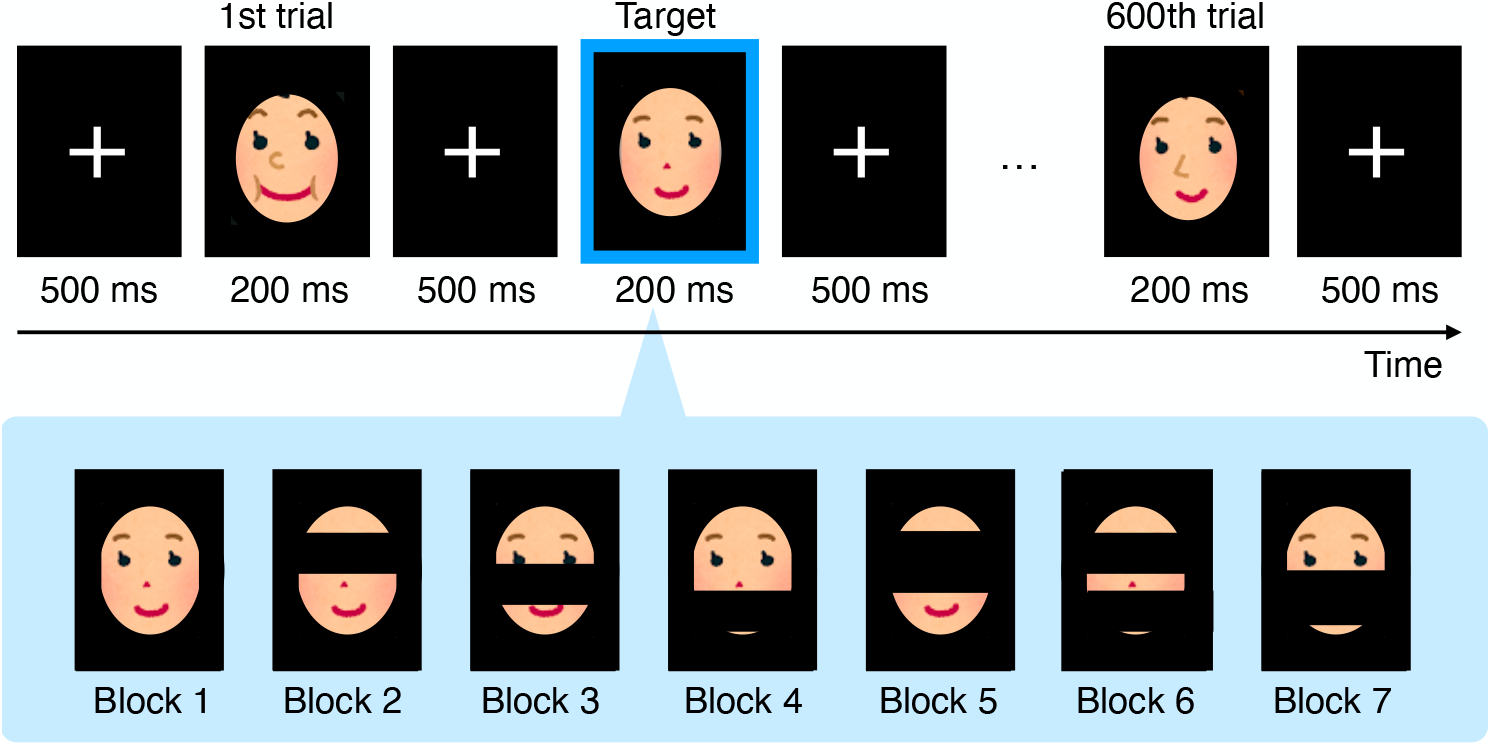
Experimental flow of the main tasks, consisting of seven blocks. In each block, only one type of face stimulus condition was shown 600 times; 100 times per face image type.

### Data Acquisition

The participants sit at a distance of 60 cm from the Monitor (EIZO, FlexScan EV2451, 23.8 inches, 1920 × 1080 pixels) of the eye tracker, Tobii Pro Spectrum (Tobii, Stockholm, Sweden). All participants were asked to sit in their comfortable positions but keep their heads and eyes on the center of the screen. During this experiment, we recorded two biosignals: EEG and eye-tracking. EEG data were acquired using g.USBamp amplifier (g.tec, Graz, Austria) at a sampling rate of 600 Hz. We used OpenViBE software version 2.2.0 (Renard et al, 2010), two photodiode triggers for recording, and Arduino for synchronizing the EEG and eye-tracking recorders. Fourteen EEG electrodes were placed on the scalps in accordance with the international 10–20 system (Fp1, Fp2, F7, F3, F4, F8, C3, Cz, C4, P3, Pz, P4, O1, and O2), with the reference electrode in the Fz position and the ground electrode on the left earlobe (A1). The eye-tracking data were recorded using a screen-based eye tracker, Tobii Pro Spectrum (Tobii, Stockholm, Sweden), at a 300 Hz sampling rate, using Tobii Pro Lab software version 1.194. For each recording, the participants were asked to perform calibration of all fixation points on the screen before the task started. The EEG and eye-tracking data were synchronized using photodiodes. The two photodiodes were used to detect the brightness changes on the screen and send the voltage to Arduino DUE. The Arduino control program, written using Arduino software version 1.8.20, received the voltage changes and sent the trigger to an OpenViBE software communication program written with Python version 3.5.8. This program tags the event and sends the stimulation to OpenViBE software.

### Data Analysis

#### Electroencephalogram (EEG)

The recorded EEG data were processed using MNE-python package 1.3.0 (Gramfort et al, 2013) on Python version 3.5.8. We first filtered with a 0.5 to 30 Hz bandpass filter to remove the low-frequency and 50 Hz noise and downsampled the data to 300 Hz to match the sampling frequency with the eye-tracking. The acquired EEG was then segmented into 600 epochs with a duration from −100 to 700 msec, having a duration of −100 to 0 msec as the baseline. The data then underwent baseline correction using the mean amplitude of the baseline period, and the epochs with the peak-to-peak amplitude over 100 *μ*V were rejected to obtain a clean data set with the least noise contamination. At this stage, we were left with 80 ± 3 epochs per image.

The clean data were averaged across the electrode group according to the brain lobe (Valiulis, 2014) shown in Figure 2. The data of all the participants are grand averaged to obtain the ERP. To test our hypothesis and investigate how partial-face cognition affected ERP compared to full-face cognition, we performed a statistical test using a paired t-test on the largest amplitude of the ERP components in the given duration (N170 at 100 to 200 msec, P3a at 200 to 250 msec, and P3b at 250 to 550 msec after the stimulus onset using JASP version 0.16.4 (JASP Team, 2023). We also tested the latency of the largest amplitude to investigate the delay of the ERP components among all the partial face cognition conditions. We applied a one-tailed dependent t-test for each amplitude and each latency of the ERP components to test that the participants performed better in full face cognition task and recognized faces using holistic processing and vice-versa to test that the partial face cognition could be used to enhance the ability of identification faces when compare full face.

**Fig. 2.**
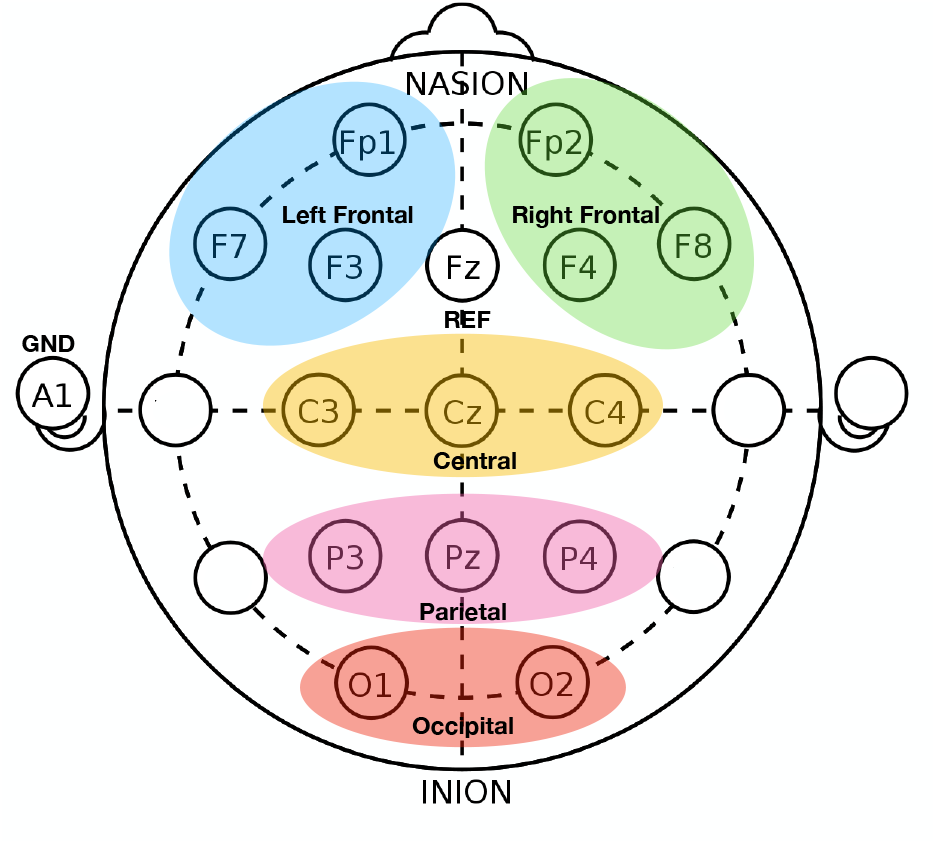
Electrode group, where blue color represents the left frontal area, green color represents the right frontal area, orange color represents the central area, pink color represents the parietal area, and red color represents the occipital area of EEG electrode position

### Eye-tracking data

We used the analysis function in Tobii Pro Lab software version 1.194 to process the eye-tracking data. This software created the heat map based on the absolute count, gaze point, and total fixation duration in the target face’s area of interest (AOI). The AOIs of the face images were marked by the three polygons for eyes, nose, and mouth components so that they covered an equal area of the face proportion, as shown in Figure 3. The time per media (200 msec) was used as the time of interest (TOI); therefore, the total duration of fixation was calculated by taking the mean of the total duration of fixation of each media (each face image type). We also calculated the total duration of fixation of the inter-stimulus interval (ISI) to investigate whether the participants kept their eyes on the fixation during the ISI by taking the mean of all the time duration the eyes fixed on the fixation during ISI.

**Fig. 3.**
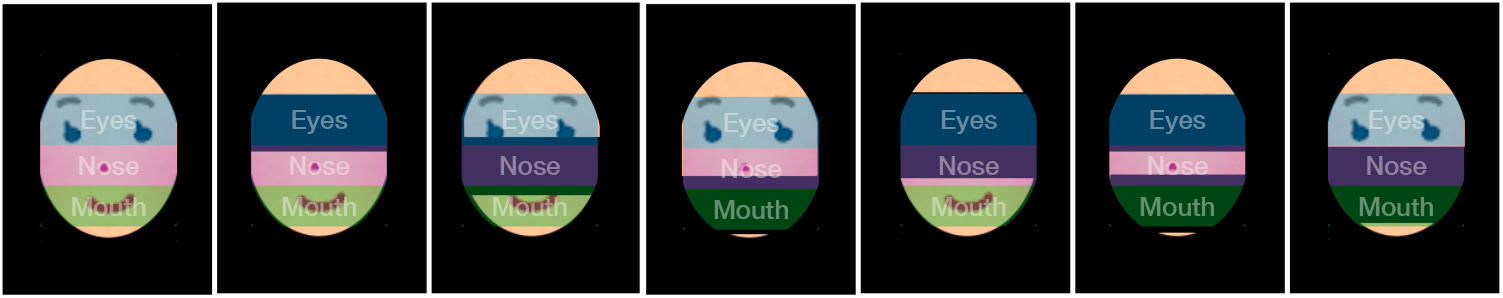
The Area of Interest (AOI) used to process the eye-tracking data

### Canonical Correlation Analysis based on EEG and Eye-Tracking

To investigate the face cognition mechanism among all the tasks, we performed canonical correlation analysis (Witten et al, 2009; Mai and Zhang, 2019) based on the total fixation duration in four AOIs (eyes, nose, mouth, and fixation) and the amplitude and latency of P3a and P3b subcomponents using Python and the sparsecca package (Teekuningas, 2019).

## 3 Results

### Event-Related Potentials (ERPs)

Figure 4 shows the ERP component changes and eye-tracking results in the cognition tasks when the full face, partial face with eyes covered, partial face with nose covered, and partial face with nose and mouth covered were used as a stimulus. The EEG results shown on the left of Figure 4 were obtained from grand averaging the EEG data of the electrode position located in the frontal and parietal area, and the heat map and gaze order of the eye-tracking data was shown on the right. From Figure 4, we could observe two ERP components, N170 found in the frontal area, and P300; P3a found in the frontal area and P3b found in the parietal area. We then performed a one-tailed dependent t-test for each peak and latency.

**Fig. 4.**
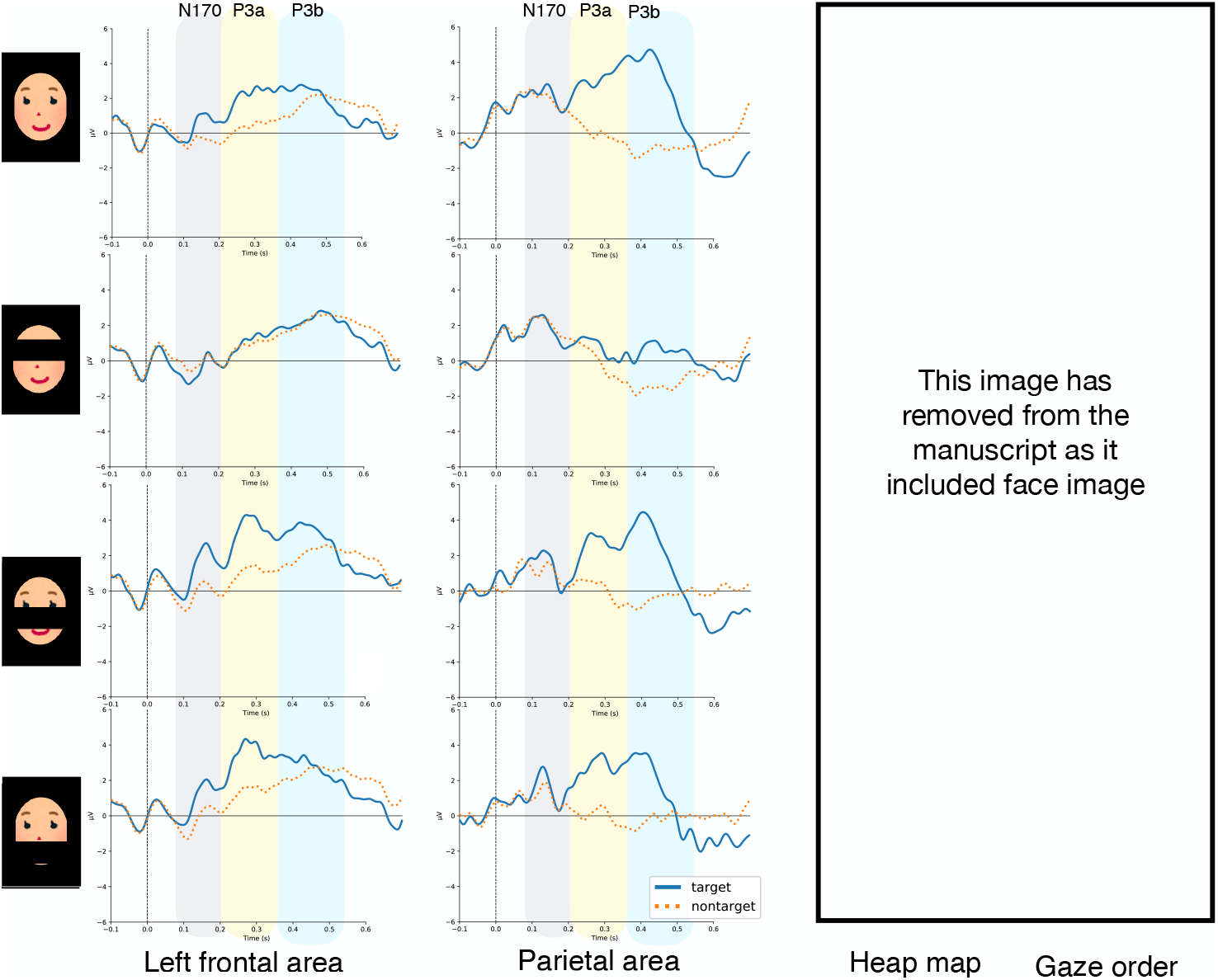
Examples of grand average ERP (left) and eye-tracking heat map and gaze order (right) of full face, partial face with eyes covered and partial face with nose covered, and partial face with nose and mouth covered condition, respectively. The grey, yellow, and blue highlighted areas on the left figure represent the latency of N170, P3a, and P3b. The thick blue line represents the ERP of the target, while the dotted orange line represents the non-target. The range of the heat map was set based on the absolute count, where the green color represents the least count, and the red color represents the most count. The color variation in the gaze order result represents each participant with a different color with 100 gazes each.

As a result, we observed that the amplitude of N170 in full face cognition was significantly larger than in the partial face with the eyes covered cognition (*t*(17) = 2.064, *p <* 0.05, *d* = 0.486, see Supplementary Table 4) as shown in Figure 5(a), but could not observe any significant latency differences between the full face and partial face cognition in N170 (see Supplementary Tables 1, 2, and 5 for more details). When we looked at the shape of the P3a component when the participants recognized the target face in Figure 4, we could not see a clear peak in the full face cognition condition as how we saw in the previous experiment (Chanpornpakdi and Tanaka, 2023b), but when we compared the target P3a peak amplitude to the amplitude of the P3a evoked by the non-target faces, the amplitude evoked by the target face was comparatively larger. In a comparison of the ERPs evoked in the full face cognition with the partial face cognition, we could clearly see that when the face with the eye component covered was used as a stimulus, both P3a and P3b peaks became significantly smaller (when only eyes were covered: *t*(17) = 2.112, *p <* 0.05, *d* = 0.498 for P3a (see Supplementary Table 10) and *t*(17) = 3.379, *p <* 0.005, *d* = 0.796 for P3b (see Supplementary Table 16); when the eyes and nose were covered: *t*(17) = 3.997, *p <* 0.001, *d* = 0.942 for P3b (see Supplementary Table 16); when eyes and mouth were covered: *t*(17) = 5.820, *p <* 0.001, *d* = 1.372 (see Supplementary Table 16)) as shown in Figures 5 (b) and 5(c). Moreover, we also found that the amplitude of P3b of the full face cognition became larger when compared to the partial face cognition with mouth covered (*t*(17) = 2.919, *p <* 0.01, *d* = 0.688, see Supplementary Table 16) as shown in Figure 5 (c). In addition, we also found that the P3a peak observed in the face cognition when the nose was covered was larger than that of the full face cognition (*t*(17) = −1.943, *p <* 0.05, *d* = −0.458, see Supplementary Table 11) as shown in Figure 5 (b). However, we could not notice any significant differences in the latency when comparing the P3a and P3b peaks evoked in full face and partial face cognition (see Supplementary Tables 7, 8, 13, and 14).

**Table 1.**
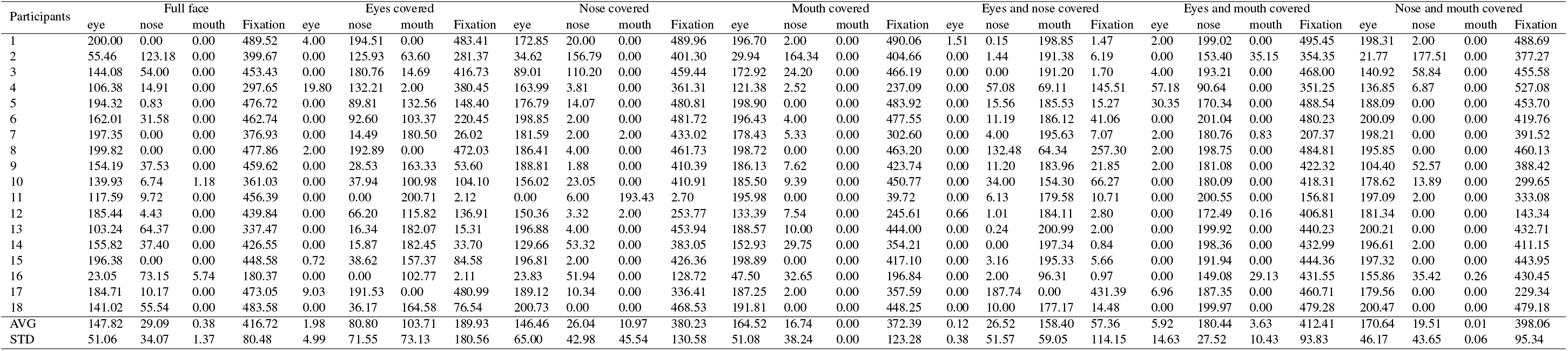
The total duration of fixation of each AOIs in all face cognition tasks.

**Fig. 5.**
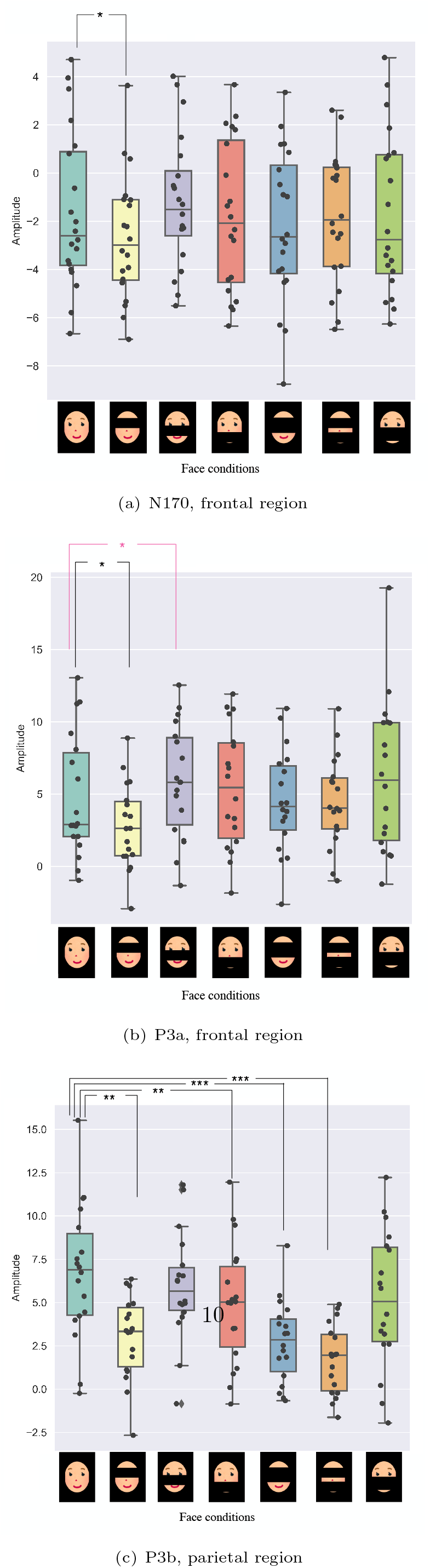
ERP amplitude comparison for each condition. (*n* = 18, *p <* 0.05∗, *p <* 0.01∗∗, *p <* 001∗∗∗) Black color indicates that the ERP amplitude of full-face cognition was larger than that of partial-face cognition, whereas pink color indicates the opposite.

### Eye-Tracking

From the middle figure in Figure 4, we could see from the heat map results that although some other components were presented along with the eyes, most eye gazes were focused on the eyes the most. The gaze order of the eye-tracking data on the right of Figure 4 shows that although the majority of the data were on the eye component, one participant focused on the mouth rather than the eyes when the face with nose covered were shown as a stimulus.

Figure 6 and Table 3 show the total duration of fixation in the AOIs in all the face cognition tasks. The total duration of fixation of AOIs was extremely low when that area of the face was covered. The eye-tracking results show that the total duration of fixation of AOIs in the full face cognition, partial-face cognition with the nose covered, and partial-face cognition with the nose the mouth covered were similar to one another, and the fixation duration in the eye AOI for the longest period among all components. In addition, we also observed that the eye fixation duration of the task using the face with the mouth covered showed similar results, too. The longest fixation duration in the eyes AOI area with the average duration of 147.82 ± 51.06 msec for the full face condition, 146.46 ± 65.00 msec for the face with nose covered condition, 164.51 ± 16.74 msec for the face with mouth covered, and 170.64 ± 46.17 msec for the face with nose and mouth covered condition (equivalent to mask condition). Among these four conditions, we noticed that when the nose and mouth were invisible, the duration of fixation at the eyes component became the longest.

**Fig. 6.**
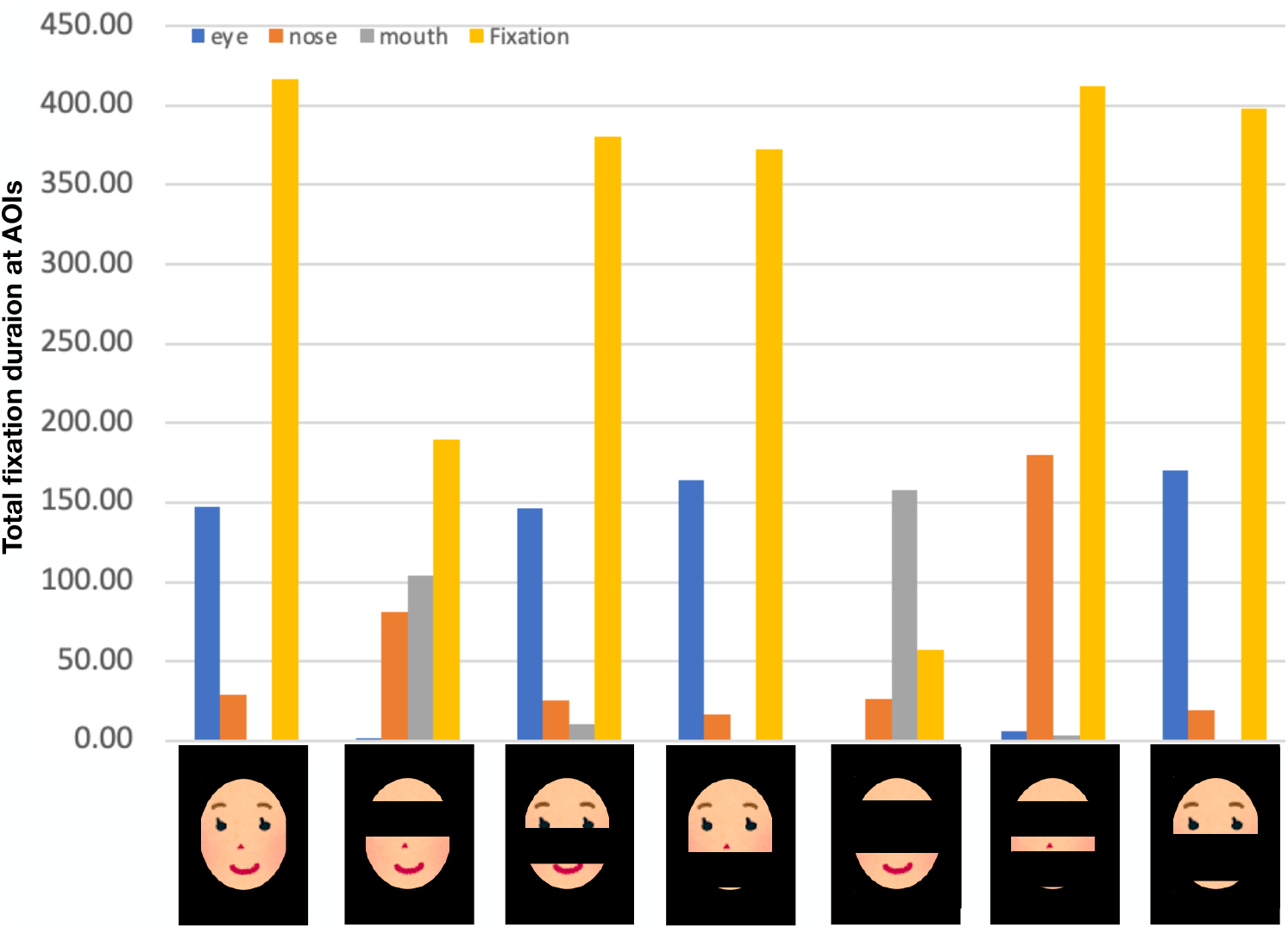
The average total fixation duration at eyes, nose, mouth, and fixation AOIs in all tasks.

From Table 3, few participants gathered and kept their eyes fixed on the nose area although the eyes were presented (full face cognition task: two participants; partial face cognition task in which the eye component was visible: three participants in the face with nose covered condition, one participant in the face with mouth covered condition, and one participant in the face with nose and mouth covered condition). The average total fixation duration of all participants looking at the nose was 29.09 ± 34.07 for full face, 26.04 ± 49.54 for the face with nose covered, 16.74 ± 38.24 for the face with mouth covered, and 19.51 ± 43.65 for the face with nose and mouth covered. We also spotted that one participant focused on the mouth rather than the eyes, which most participants did when the face with the nose covered was used as a stimulus.

### Canonical Correlation Analysis based on EEG and Eye-Tracking

To investigate how each task of face cognition is correlated to each other based on the EEG and eye-tracking data, we calculated the canonical correlation of all tasks using the total duration fixation of each AOI (eyes, nose, mouth, and fixation) from Figure 6 and Table 3, and the amplitude and latency of P3a and P3b peaks. We then plotted the confusion matrix for the canonical correlation coefficient. Figure 7 shows the canonical correlation between each face cognition task calculated using EEG and eye-tracking data. From this figure, we can see that all the tasks are positively correlated. The cognition of the full face was highly correlated with the cognition of the face with mouth-covered cognition (*R*_*c*_ = 0.88) and cognition of the face with nose and mouth covered (*R*_*c*_ = 0.93). We also observed that the cognition of the face with mouth covered and the face with nose and mouth covered is also closely associated (*R*_*c*_ = 0.93).

**Fig. 7.**
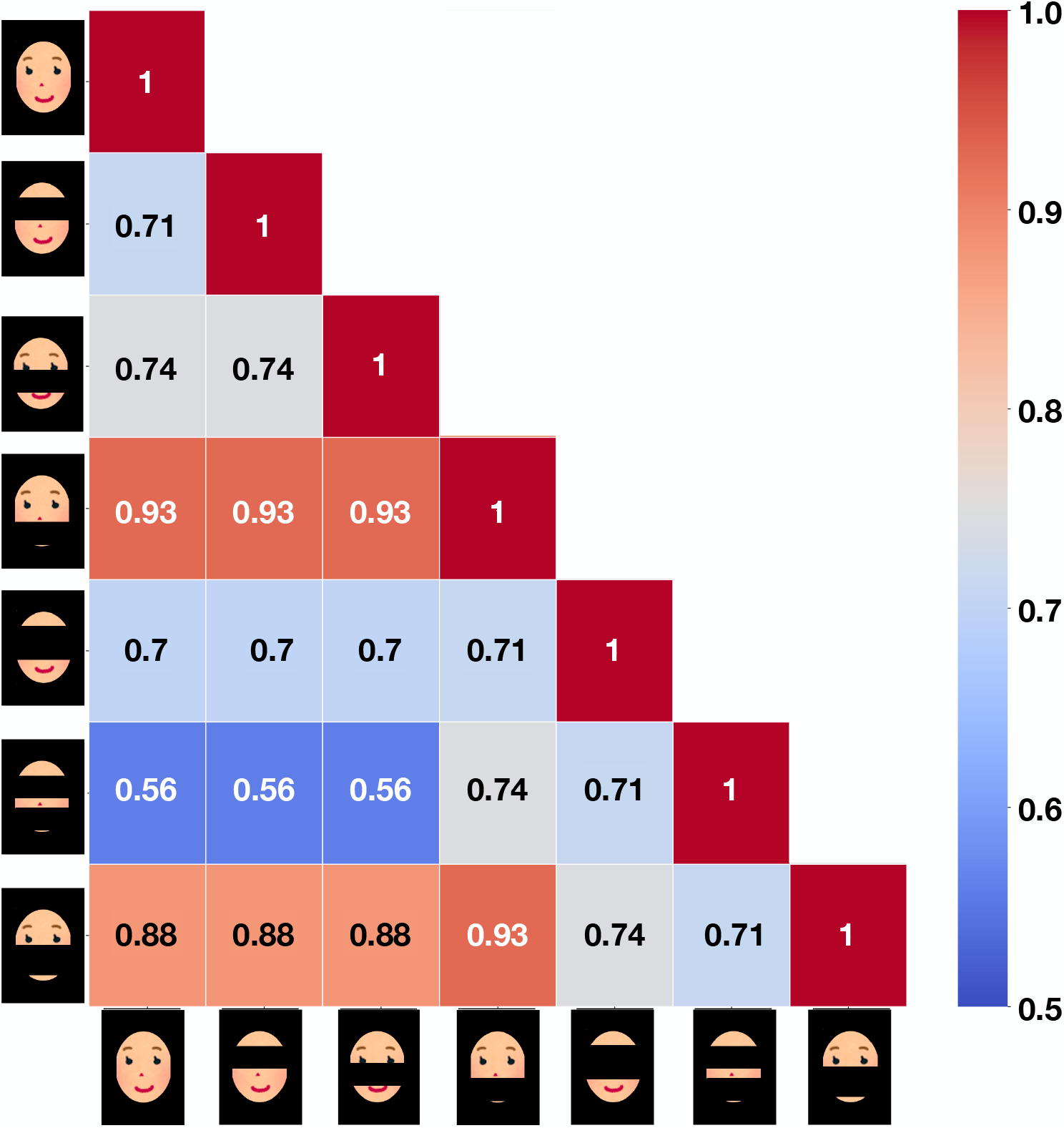
The canonical correlation of each face cognition task using EEG and eye-tracking data.

## 4 Discussion

In this study, we investigated the partial face mechanism using EEG to examine the evoked ERP components changes across the face cognition condition and eye-tracking to examine which area of the face the participants focus on. We found two ERP components (N170 and P300; P3a and P3b) evoked as brain responses in face cognition. The amplitudes and latencies of N170 showed no significant difference among all face conditions, whereas the amplitude of P3a and P3b components became smaller, and the latency of P3a and P3b components got extended when compared to the amplitude evoked by the full face cognition. In addition, we also found that the amplitude of P3a in the condition when the participants recognize faces with the nose cover became larger than the full face cognition. For eye-tracking data, we found that most participants fixed their eyes on the eye area whenever the eyes were presented. We also observed similar participants during the full face, face with nose covered, face with mouth covered, and face with nose and mouth covered cognition tasks. Using the two data, we calculated the canonical correlation among the face cognition condition and found that the full face cognition and partial face with the mouth covered and the partial face with the nose and mouth covered cognition were highly correlated.

### Event-Related Potential (ERP)

From our experiment, we have found two components of ERPs, N170 and P300. The N170 and P3a peak was found in the frontal area of the brain, whereas P3b was found in the parietal area of the brain. N170 is one of the face characteristic ERPs that can be induced, especially during the cognition task when the target face was shown among the other objects (Joyce and Rossion, 2005; Eimer, 2011; Cai et al, 2013). Some studies found significant differences in either latency or amplitude or both of them during partial face cognition (Nemrodov et al, 2014; Rousselet et al, 2014; Ince et al, 2016; Żochowska et al, 2022) whereas we did not find any significant differences in either of the results. The explanation for such a difference in results might be that more factors than face components affect the cognition process, such as presentation time and how the participants were asked to respond during the task.

The reason why we found the subcomponents of P300 peak larger at different parts of the brain is that P3a is known for the focal attention, which functional area is located in the frontal lobe, whereas P3b is responsible for the target cognition (Bressler and Ding, 2006; Polich and Criado, 2006; Polich, 2007). In addition, the left inferior frontal cortex is the brain area that processes language as well as memory retrieval (Lundstrom et al, 2005). The P3a evoked might also relate to memory retrieval as the participants tried to recall the face they remembered and tried to retrieve the moment when they were remembering the face. This activity may activate the precuneus in the frontal cortex, which is sensitive to the quantity and quality of the information retrieved (Rugg et al, 1998; Nyberg et al, 2000; Lundstrom et al, 2003, 2005).

The frontal cortex also functions in the visual working memory (Courtney et al, 1998; Chai et al, 2018). Vogel and Machizawa (2004) studied the capacity of the human visual working memory capacity and found that as the number of items increased, the ERP response also became larger, but at some point, it reached a limit. They interpreted that individuals have different memory capacities and found that each has a capacity with an average of about 2.8 items. When we look at the face features, the face contains four components: two eyes (consider the eyebrow as a part of the eye component), one nose, and one mouth. Covering the nose could reduce the number of components into three components almost equal to the capacity that Vogel and Machizawa (2004) have found. This could explain why recognizing the face with the nose covered results in a larger P3a component, not P3b. When we covered both the nose and mouth, we had fewer components, but we could still see a similar level of P3a and P3b peaks with the condition of the partial face with the nose covered. This might be because the eye component is the most important part of face cognition, as we can see when we covered the eye component and asked the participant to recognize the face. Both P3a and P3b dropped significantly. One more reason behind that could be due to neuroplasticity. As we have been living with the mask on for many years, people could learn and get used to recognizing faces with the nose and mouth covered as time passed. This makes us recognize faces with less difficulty and results in a similar level of P3a and P3b even though we have only a few components (only two but important components).

### Eye-Tracking

To investigate that the eye component is the most informative feature in face cognition, we have recorded the eye-tracking data along with EEG and found that most participants focused on the eyes the most and fixed their gaze on the eye area. We assumed that the total duration of fixation of AOIs in the full face cognition, partial-face cognition with the nose covered, and partial-face cognition with the nose and mouth covered would be similar to one another, and the fixation duration in the eye AOI would be the longest period among all components, as they showed in the similar level of accuracy in the classification model in the target classification results based on ERP in our previous study (Chanpornpakdi and Tanaka, 2023b). The eye-tracking results coincide with our assumption. In addition, we also observed that the eye fixation duration of the task using the face with the mouth covered showed similar results to our assumption. These results coincided with our ERP results that when the eye component was covered, the participants lost some ability to recognize faces and confirmed that people use eye information the most, and it plays an important role in face cognition (Nguyen and Pezdek, 2017; Royer et al, 2018). We also found that some participants still focused on the nose, the fixation position, although the nose was invisible. From these results, we can interpret that people can still take the face as a whole and recognize faces using holistic face processing even though some part of the face is missing. The reason might be that we still presented the eyes and have the visible components in the correct position as people can still recognize faces using holistic processing when the facial features are in the correct position rather than selecting parts of them and identifying the face (de Haas et al, 2016; Poltoratski et al, 2021).

Contradictory to the majority, we saw one participant looking at the mouth rather than the eyes when the face with the nose covered was used as a stimulus. There could be two possibilities as to why this person focused on the mouth. This person could have a larger visual field coverage of face-selective regions so that it included the eye area even though the person’s fixation was on the mouth (Poltoratski et al, 2021). Another possibility is that we have selected the target image with the nose and mouth shape that stood out from the rest of the images. This could make the target look different from the Asian face and cause the person to identify the ethnic race rather than the facial identity as people tend to focus on the eyes when identifying a face identity, whereas looking at the mouth when identifying the ethnic (Chakravarthula et al, 2021). From our results, we could interpret that partial face cognition might rely on holistic face processing if we have the important features of the face in the right position and structural face processing depending on the individual. This could give the potential to use partial face cognition to enhance the face cognition task.

### Canonical Correlation Analysis based on EEG and Eye-Tracking

From our canonical correlation analysis results, we observed that the full face cognition, the face with the mouth covered, and the face with nose and mouth covered have strong correlations. The reason could be that the face with the nose and mouth covered was the partial face condition that resembles the surgical mask the most. Moreover, The face with the mouth covered is similar to nose commando (Wolff, 2020), the condition when people wear a half mask with the nose exposed. This indicates that people can recognize the partial face with mouth covered and the partial face with nose and mouth covered comparably with the full face cognition based on the EEG and eye-tracking data. We can interpret that the cognition mechanism of the masked face and the full face are similar; hence, people can recognize faces correctly even though some features of the face are covered under the mask.

### Limitation and Solution

Regarding the stimulus design, because we used color images and did not remove the remarkable facial features such as moles, this could distract the participants and attract more attention than the major components we aimed to investigate. In addition, the presentation time was 200 msec, which is considered shorter than the suitable time presentation for face cognition that Lees et al (2018) suggested. Another limitation of our experiment was that we had a fixed duration for the interstimulus interval (ISI), which could lead the participant to anticipatory effects. To prevent such an effect, we should jitter the ISI randomly to reduce the cue of when the stimulus will be presented and obtain the brain response purely based on the cognitive function.

Moreover, even though we have reduced the task to lighten the work burden during the experiment, we have increased the number of trials from 20 trials/image to 100 trials/image in order to obtain clean EEG data, the workload was still considered to be heavy and could cause the fatigue and reduce the ERP responses. To solve such a problem, we should divide each face cognition condition block into smaller sessions so the participants can work on face recognition on a small batch of face images and get some time to rest between sessions. We also found that the latencies of evoked ERPs were slower than those we found from the experiment in our previous study (Chanpornpakdi and Tanaka, 2023b). The reason could be that we created our own self-made trigger using photodiodes, resistors, and Arduino to synchronize the EEG and eye-tracking data.

To synchronize the data, we attached the photodiodes to the screen to detect the light changes on the screen, used Arduino software to control Arduino and send the trigger signal to OpenViBE software when there were changes on the screen, then we programmed OpenViBE software to send the external triggers signal to the EEG amplifier using python. All these processes could cause time in accuracy in the trigger due to the frame refresh rate and data transfer, such as delay settings in the controller and external trigger programs. To solve the problem, we need to identify the exact delay by measuring the delay caused by both the Arduino control program and the OpenViBE communication program. We should also consider using the trigger synchronization system sold on the market so that we can minimize time inaccuracy and improve our data quality.

### Future Work

In the future, we plan to analyze our eye-tracking data in more detail to investigate the correlation between the eye gaze area and EEG response and implement a hidden Markov Model (Rabiner and Juang, 1986; Rabiner, 1989; Chuk et al, 2014) to identify how the eye gaze develops during the experiment and the interaction between the AOIs. For future applications of EEG-based RSVP, we intend to apply these results to create a lie detector using a partial face image. We believe that partial face image stimulus could take the revealing concealed face information task to the next level due to the absence of some face features. This stimulus can make the cognition task of the concealed target face become difficult to deceive. Another application could be a marketing analyzer using EEGs to investigate customer preference in selecting products.

## Supplementary information

The details of the statistical results are available in the Supplementary Materials.

## Data availability statement

Physiological signals (EEG and eye-tracking) were recorded for this study. The experimental protocol was approved by the Ethics Committee, but the committee’s decision did not permit the publication of the data.

The software libraries used in the analysis are described in the Methods section.

## Acknowledgments

This work was supported in part by JST Moonshot R&D Grant Number JPMJMS2237.

