## Supplementary Table for "Partial Face Visibility and Facial Cognition: An Electroencephalography and Eye-Tracking Investigation"

TABLE 1: Paired Samples T-Test (N170 latency of full face > partial face at frontal area)

| Measure 1 | | Measure 2 | t | df | p | Mean Difference | SE Difference | Cohen's d | SE Cohen's d |
|---|---|---|---|---|---|---|---|---|---|
| Full face (Latency) | - | Eyes (Latency) | −0.105 | 17 | 0.541 | −0.001 | 0.011 | −0.025 | 0.299 |
| | - | Nose (Latency) | 1.655 | 17 | 0.058 | 0.012 | 0.007 | 0.390 | 0.209 |
| | - | Mouth (Latency) | −0.468 | 17 | 0.677 | −0.002 | 0.005 | −0.110 | 0.124 |
| | - | Eyes and nose (Latency) | 1.014 | 17 | 0.162 | 0.010 | 0.010 | 0.239 | 0.300 |
| | - | Eyes and mouth (Latency) | 0.158 | 17 | 0.438 | 0.001 | 0.008 | 0.037 | 0.237 |
| | - | Nose and mouth (Latency) | 1.562 | 17 | 0.068 | 0.009 | 0.006 | 0.368 | 0.156 |

TABLE 2: Paired Samples T-Test (N170 latency of full face < partial face at frontal area)

| Measure 1 | | Measure 2 | t | df | p | Mean Difference | SE Difference | Cohen's d | SE Cohen's d |
|---|---|---|---|---|---|---|---|---|---|
| Full face (Latency) | - | Eyes (Latency) | −0.105 | 17 | 0.459 | −0.001 | 0.011 | −0.025 | 0.299 |
| | - | Nose (Latency) | 1.655 | 17 | 0.942 | 0.012 | 0.007 | 0.390 | 0.209 |
| | - | Mouth (Latency) | −0.468 | 17 | 0.323 | −0.002 | 0.005 | −0.110 | 0.124 |
| | - | Eyes and nose (Latency) | 1.014 | 17 | 0.838 | 0.010 | 0.010 | 0.239 | 0.300 |
| | - | Eyes and mouth (Latency) | 0.158 | 17 | 0.562 | 0.001 | 0.008 | 0.037 | 0.237 |
| | - | Nose and mouth (Latency) | 1.562 | 17 | 0.932 | 0.009 | 0.006 | 0.368 | 0.156 |

TABLE 3: Descriptives (N170 latency)

| | N | Mean | SD | SE | Coefficient of variation |
|---|---|---|---|---|---|
| Full face (Latency) | 18 | 0.134 | 0.039 | 0.009 | 0.288 |
| Eyes (Latency) | 18 | 0.135 | 0.032 | 0.007 | 0.234 |
| Nose (Latency) | 18 | 0.122 | 0.035 | 0.008 | 0.290 |
| Mouth (Latency) | 18 | 0.136 | 0.043 | 0.010 | 0.315 |
| Eyes and nose (Latency) | 18 | 0.124 | 0.022 | 0.005 | 0.179 |
| Eyes and mouth (Latency) | 18 | 0.133 | 0.029 | 0.007 | 0.216 |
| Nose and mouth (Latency) | 18 | 0.125 | 0.037 | 0.009 | 0.292 |

TABLE 4: Paired Samples T-Test (N170 amplitude of full face > partial face at frontal area)

| Measure 1 | | Measure 2 | t | df | p | Mean Difference | SE Difference | Cohen's d | SE Cohen's d |
|---|---|---|---|---|---|---|---|---|---|
| Full face (Amplitude) | - | Eyes (Amplitude) | 2.064 | 17 | 0.027 | 1.088 | 0.527 | 0.486 | 0.182 |
| | - | Nose (Amplitude) | −1.109 | 17 | 0.859 | −0.445 | 0.401 | −0.261 | 0.128 |
| | - | Mouth (Amplitude) | 0.261 | 17 | 0.399 | 0.122 | 0.468 | 0.062 | 0.147 |
| | - | Eyes and nose (Amplitude) | 1.074 | 17 | 0.149 | 0.580 | 0.540 | 0.253 | 0.171 |
| | - | Eyes and mouth (Amplitude) | 0.353 | 17 | 0.364 | 0.185 | 0.525 | 0.083 | 0.172 |
| | - | Nose and mouth (Amplitude) | 0.118 | 17 | 0.454 | 0.059 | 0.503 | 0.028 | 0.152 |

TABLE 5: Paired Samples T-Test (N170 amplitude of full face < partial face at frontal area)

| Measure 1 | | Measure 2 | t | df | p | Mean Difference | SE Difference | Cohen's d | SE Cohen's d |
|---|---|---|---|---|---|---|---|---|---|
| Full face (Amplitude) | - | Eyes (Amplitude) | 2.064 | 17 | 0.973 | 1.088 | 0.527 | 0.486 | 0.182 |
| | - | Nose (Amplitude) | −1.109 | 17 | 0.141 | −0.445 | 0.401 | −0.261 | 0.128 |
| | - | Mouth (Amplitude) | 0.261 | 17 | 0.601 | 0.122 | 0.468 | 0.062 | 0.147 |
| | - | Eyes and nose (Amplitude) | 1.074 | 17 | 0.851 | 0.580 | 0.540 | 0.253 | 0.171 |
| | - | Eyes and mouth (Amplitude) | 0.353 | 17 | 0.636 | 0.185 | 0.525 | 0.083 | 0.172 |
| | - | Nose and mouth (Amplitude) | 0.118 | 17 | 0.546 | 0.059 | 0.503 | 0.028 | 0.152 |

TABLE 6: Descriptives (N170 amplitude)

| | N | Mean | SD | SE | Coefficient of variation |
|---|---|---|---|---|---|
| Full face (Amplitude) | 18 | −1.278 | 3.281 | 0.773 | −2.567 |
| Eyes (Amplitude) | 18 | −2.366 | 2.483 | 0.585 | −1.050 |
| Nose (Amplitude) | 18 | −0.833 | 2.631 | 0.620 | −3.157 |
| Mouth (Amplitude) | 18 | −1.400 | 3.078 | 0.725 | −2.198 |
| Eyes and nose (Amplitude) | 18 | −1.858 | 3.138 | 0.740 | −1.689 |
| Eyes and mouth (Amplitude) | 18 | −1.463 | 2.567 | 0.605 | −1.755 |
| Nose and mouth (Amplitude) | 18 | −1.337 | 3.348 | 0.789 | −2.504 |

TABLE 7: Paired Samples T-Test (P3a latency of full face > partial face at frontal area)

| Measure 1 | | Measure 2 | t | df | p | Mean Difference | SE Difference | Cohen's d | SE Cohen's d |
|---|---|---|---|---|---|---|---|---|---|
| Full face (Latency) | - | Eyes (Latency) | −3.023 | 17 | 0.996 | −0.019 | 0.006 | −0.713 | 0.180 |
| | - | Nose (Latency) | −2.145 | 17 | 0.977 | −0.008 | 0.004 | −0.506 | 0.100 |
| | - | Mouth (Latency) | −1.182 | 17 | 0.873 | −0.007 | 0.006 | −0.279 | 0.150 |
| | - | Eyes and nose (Latency) | −2.473 | 17 | 0.988 | −0.024 | 0.010 | −0.583 | 0.248 |
| | - | Eyes and mouth (Latency) | −0.500 | 17 | 0.688 | −0.005 | 0.010 | −0.118 | 0.251 |
| | - | Nose and mouth (Latency) | −0.107 | 17 | 0.542 | $-7.407 \times 10^{-4}$ | 0.007 | −0.025 | 0.187 |

TABLE 8: Paired Samples T-Test (P3a latency of full face < partial face at frontal area)

| Measure 1 | | Measure 2 | t | df | p | Mean Difference | SE Difference | Cohen's d | SE Cohen's d |
|---|---|---|---|---|---|---|---|---|---|
| Full face (Latency) | - | Eyes (Latency) | −3.023 | 17 | 0.996 | −0.019 | 0.006 | −0.713 | 0.180 |
| | - | Nose (Latency) | −2.145 | 17 | 0.977 | −0.008 | 0.004 | −0.506 | 0.100 |
| | - | Mouth (Latency) | −1.182 | 17 | 0.873 | −0.007 | 0.006 | −0.279 | 0.150 |
| | - | Eyes and nose (Latency) | −2.473 | 17 | 0.988 | −0.024 | 0.010 | −0.583 | 0.248 |
| | - | Eyes and mouth (Latency) | −0.500 | 17 | 0.688 | −0.005 | 0.010 | −0.118 | 0.251 |
| | - | Nose and mouth (Latency) | −0.107 | 17 | 0.542 | $-7.407 \times 10^{-4}$ | 0.007 | −0.025 | 0.187 |

TABLE 9: Descriptives (P3a latency)

| | N | Mean | SD | SE | Coefficient of variation |
|---|---|---|---|---|---|
| Full face (Latency) | 18 | 0.292 | 0.038 | 0.009 | 0.132 |
| Eyes (Latency) | 18 | 0.310 | 0.038 | 0.009 | 0.124 |
| Nose (Latency) | 18 | 0.299 | 0.039 | 0.009 | 0.129 |
| Mouth (Latency) | 18 | 0.299 | 0.040 | 0.009 | 0.135 |
| Eyes and nose (Latency) | 18 | 0.315 | 0.044 | 0.010 | 0.139 |
| Eyes and mouth (Latency) | 18 | 0.297 | 0.042 | 0.010 | 0.140 |
| Nose and mouth (Latency) | 18 | 0.292 | 0.035 | 0.008 | 0.121 |

TABLE 10: Paired Samples T-Test (P3a amplitude of full face > partial face at frontal area)

| Measure 1 | | Measure 2 | t | df | p | Mean Difference | SE Difference | Cohen's d | SE Cohen's d |
|---|---|---|---|---|---|---|---|---|---|
| Full face (Amplitude) | - | Eyes (Amplitude) | 2.112 | 17 | 0.025 | 1.985 | 0.940 | 0.498 | 0.266 |
| | - | Nose (Amplitude) | −1.943 | 17 | 0.966 | −1.223 | 0.629 | −0.458 | 0.162 |
| | - | Mouth (Amplitude) | −1.081 | 17 | 0.853 | −0.785 | 0.726 | −0.255 | 0.176 |
| | - | Eyes and nose (Amplitude) | −0.009 | 17 | 0.504 | −0.009 | 1.005 | −0.002 | 0.254 |
| | - | Eyes and mouth (Amplitude) | 0.274 | 17 | 0.394 | 0.279 | 1.018 | 0.065 | 0.269 |
| | - | Nose and mouth (Amplitude) | −1.306 | 17 | 0.896 | −1.527 | 1.169 | −0.308 | 0.251 |

TABLE 11: Paired Samples T-Test (P3a amplitude of full face < partial face at frontal area)

| Measure 1 | | Measure 2 | t | df | p | Mean Difference | SE Difference | Cohen's d | SE Cohen's d |
|---|---|---|---|---|---|---|---|---|---|
| Full face (Amplitude) | - | Eyes (Amplitude) | 2.112 | 17 | 0.975 | 1.985 | 0.940 | 0.498 | 0.266 |
| | - | Nose (Amplitude) | −1.943 | 17 | 0.034 | −1.223 | 0.629 | −0.458 | 0.162 |
| | - | Mouth (Amplitude) | −1.081 | 17 | 0.147 | −0.785 | 0.726 | −0.255 | 0.176 |
| | - | Eyes and nose (Amplitude) | −0.009 | 17 | 0.496 | −0.009 | 1.005 | −0.002 | 0.254 |
| | - | Eyes and mouth (Amplitude) | 0.274 | 17 | 0.606 | 0.279 | 1.018 | 0.065 | 0.269 |
| | - | Nose and mouth (Amplitude) | −1.306 | 17 | 0.104 | −1.527 | 1.169 | −0.308 | 0.251 |

TABLE 12: Descriptives (P3a amplitude)

| | N | Mean | SD | SE | Coefficient of variation |
|---|---|---|---|---|---|
| Full face (Amplitude) | 18 | 4.795 | 4.262 | 1.005 | 0.889 |
| Eyes (Amplitude) | 18 | 2.811 | 2.898 | 0.683 | 1.031 |
| Nose (Amplitude) | 18 | 6.019 | 3.826 | 0.902 | 0.636 |
| Mouth (Amplitude) | 18 | 5.580 | 4.111 | 0.969 | 0.737 |
| Eyes and nose (Amplitude) | 18 | 4.805 | 3.571 | 0.842 | 0.743 |
| Eyes and mouth (Amplitude) | 18 | 4.517 | 3.142 | 0.741 | 0.696 |
| Nose and mouth (Amplitude) | 18 | 6.323 | 5.163 | 1.217 | 0.817 |

TABLE 13: Paired Samples T-Test (P3b latency of full face $>$ partial face at parietal area)

| Measure 1 | | Measure 2 | t | df | p | Mean Difference | SE Difference | Cohen's d | SE Cohen's d |
|---|---|---|---|---|---|---|---|---|---|
| Full face (Latency) | - | Eyes (Latency) | −1.674 | 17 | 0.944 | −0.036 | 0.021 | −0.395 | 0.362 |
| | - | Nose (Latency) | 0.886 | 17 | 0.194 | 0.010 | 0.011 | 0.209 | 0.193 |
| | - | Mouth (Latency) | 0.585 | 17 | 0.283 | 0.009 | 0.016 | 0.138 | 0.258 |
| | - | Eyes and nose (Latency) | −2.096 | 17 | 0.974 | −0.041 | 0.019 | −0.494 | 0.293 |
| | - | Eyes and mouth (Latency) | −1.750 | 17 | 0.951 | −0.041 | 0.023 | −0.412 | 0.358 |
| | - | Nose and mouth (Latency) | 0.452 | 17 | 0.328 | 0.007 | 0.016 | 0.107 | 0.272 |

TABLE 14: Paired Samples T-Test (P3b latency of full face < partial face at parietal area)

| Measure 1 | | Measure 2 | t | df | p | Mean Difference | SE Difference | Cohen's d | SE Cohen's d |
|---|---|---|---|---|---|---|---|---|---|
| Full face (Latency) | - | Eyes (Latency) | −1.674 | 17 | 0.944 | −0.036 | 0.021 | −0.395 | 0.362 |
| | - | Nose (Latency) | 0.886 | 17 | 0.194 | 0.010 | 0.011 | 0.209 | 0.193 |
| | - | Mouth (Latency) | 0.585 | 17 | 0.283 | 0.009 | 0.016 | 0.138 | 0.258 |
| | - | Eyes and nose (Latency) | −2.096 | 17 | 0.974 | −0.041 | 0.019 | −0.494 | 0.293 |
| | - | Eyes and mouth (Latency) | −1.750 | 17 | 0.951 | −0.041 | 0.023 | −0.412 | 0.358 |
| | - | Nose and mouth (Latency) | 0.452 | 17 | 0.328 | 0.007 | 0.016 | 0.107 | 0.272 |

TABLE 15: Descriptives (P3b latency)

| | N | Mean | SD | SE | Coefficient of variation |
|---|---|---|---|---|---|
| Full face (Amplitude) | 18 | 4.795 | 4.262 | 1.005 | 0.889 |
| Eyes (Amplitude) | 18 | 2.811 | 2.898 | 0.683 | 1.031 |
| Nose (Amplitude) | 18 | 6.019 | 3.826 | 0.902 | 0.636 |
| Mouth (Amplitude) | 18 | 5.580 | 4.111 | 0.969 | 0.737 |
| Eyes and nose (Amplitude) | 18 | 4.805 | 3.571 | 0.842 | 0.743 |
| Eyes and mouth (Amplitude) | 18 | 4.517 | 3.142 | 0.741 | 0.696 |
| Nose and mouth (Amplitude) | 18 | 6.323 | 5.163 | 1.217 | 0.817 |

TABLE 16: Paired Samples T-Test (P3b amplitude of full face > partial face at parietal area)

| Measure 1 | | Measure 2 | t | df | p | Mean Difference | SE Difference | Cohen's d | SE Cohen's d |
|---|---|---|---|---|---|---|---|---|---|
| Full face (Amplitude) | - | Eyes (Amplitude) | 3.379 | 17 | 0.002 | 3.726 | 1.103 | 0.796 | 0.387 |
| | - | Nose (Amplitude) | 0.911 | 17 | 0.188 | 0.680 | 0.747 | 0.215 | 0.208 |
| | - | Mouth (Amplitude) | 2.919 | 17 | 0.005 | 1.841 | 0.631 | 0.688 | 0.186 |
| | - | Eyes and nose (Amplitude) | 3.997 | 17 | $< .001$ | 3.896 | 0.975 | 0.942 | 0.355 |
| | - | Eyes and mouth (Amplitude) | 5.820 | 17 | $< .001$ | 5.128 | 0.881 | 1.372 | 0.370 |
| | - | Nose and mouth (Amplitude) | 1.571 | 17 | 0.067 | 1.676 | 1.067 | 0.370 | 0.273 |

TABLE 17: Paired Samples T-Test (P3b amplitude of full face < partial face at parietal area)

| Measure 1 | | Measure 2 | t | df | p | Mean Difference | SE Difference | Cohen's d | SE Cohen's d |
|---|---|---|---|---|---|---|---|---|---|
| Full face (Amplitude) | - | Eyes (Amplitude) | 3.379 | 17 | 0.998 | 3.726 | 1.103 | 0.796 | 0.387 |
| | - | Nose (Amplitude) | 0.911 | 17 | 0.812 | 0.680 | 0.747 | 0.215 | 0.208 |
| | - | Mouth (Amplitude) | 2.919 | 17 | 0.995 | 1.841 | 0.631 | 0.688 | 0.186 |
| | - | Eyes and nose (Amplitude) | 3.997 | 17 | 1.000 | 3.896 | 0.975 | 0.942 | 0.355 |
| | - | Eyes and mouth (Amplitude) | 5.820 | 17 | 1.000 | 5.128 | 0.881 | 1.372 | 0.370 |
| | - | Nose and mouth (Amplitude) | 1.571 | 17 | 0.933 | 1.676 | 1.067 | 0.370 | 0.273 |

TABLE 18: Descriptives (P3b amplitude)

| | N | Mean | SD | SE | Coefficient of variation |
|---|---|---|---|---|---|
| Full face (Amplitude) | 18 | 6.710 | 3.945 | 0.930 | 0.588 |
| Eyes (Amplitude) | 18 | 2.983 | 2.420 | 0.570 | 0.811 |
| Nose (Amplitude) | 18 | 6.030 | 3.085 | 0.727 | 0.512 |
| Mouth (Amplitude) | 18 | 4.868 | 3.534 | 0.833 | 0.726 |
| Eyes and nose (Amplitude) | 18 | 2.814 | 2.352 | 0.554 | 0.836 |
| Eyes and mouth (Amplitude) | 18 | 1.582 | 2.138 | 0.504 | 1.352 |
| Nose and mouth (Amplitude) | 18 | 5.034 | 4.118 | 0.971 | 0.818 |
